## Supplementary Information for "CRISPR-mediated Multiplexed Live Cell Imaging of Nonrepetitive Genomic Loci"

Clow *et al*.

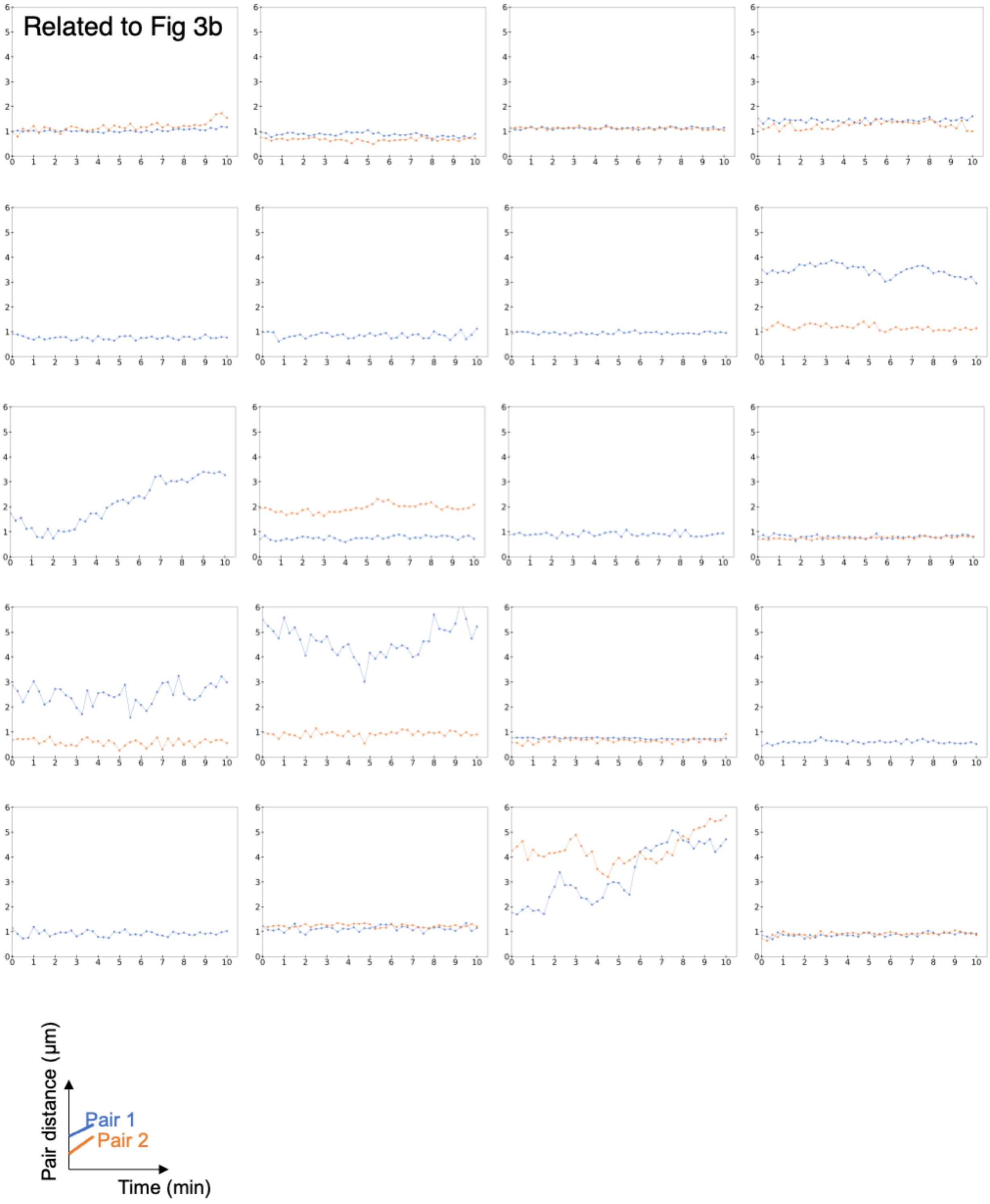


**Supplementary Fig. 1.** Pairwise 3D distances of Clover and iRFP670 foci labeling the *MASP1-BCL6* loop in 20 ARPE-19 nuclei (33 pairs) over time.

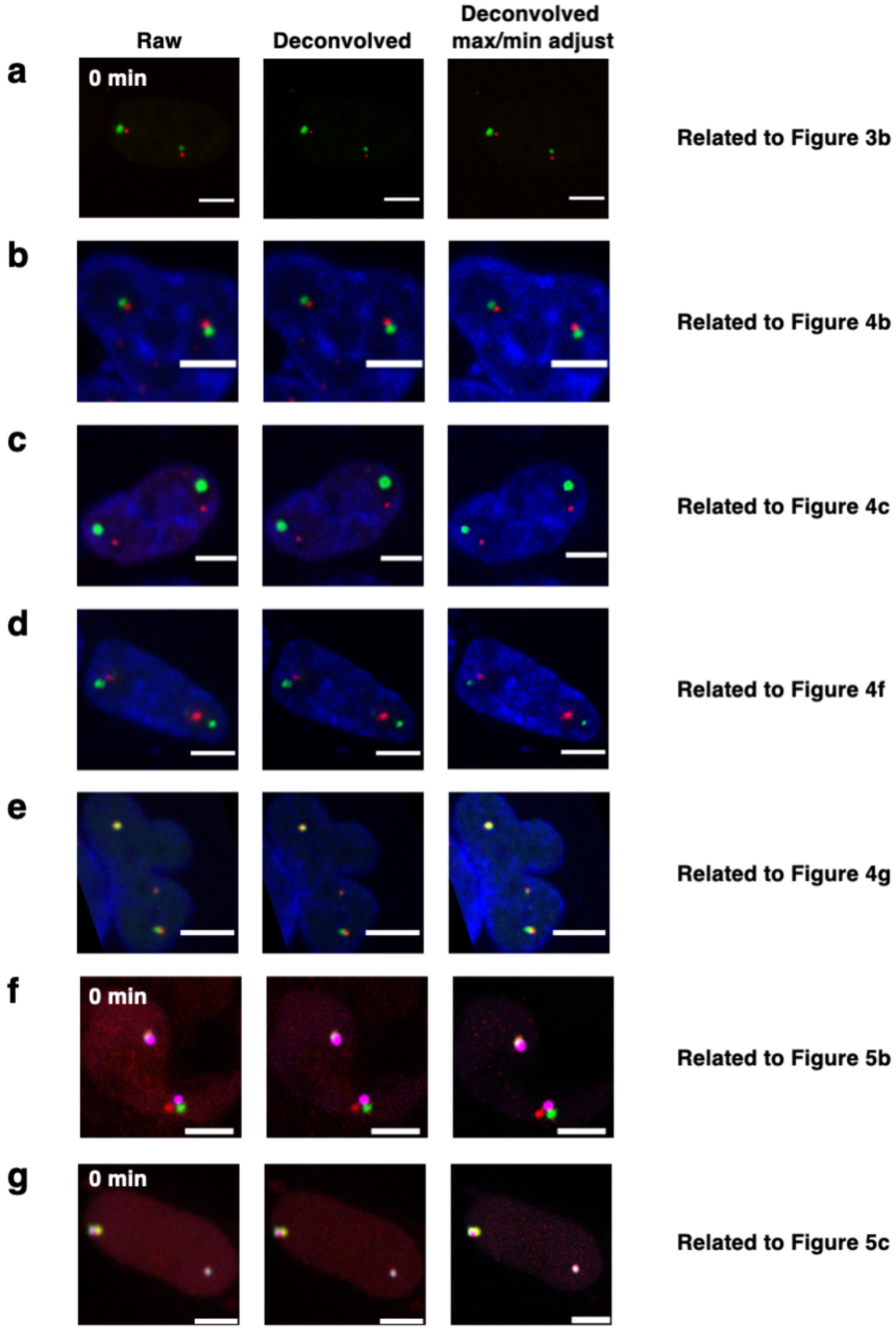


**Supplementary Fig. 2.** Example images through each step of image processing.

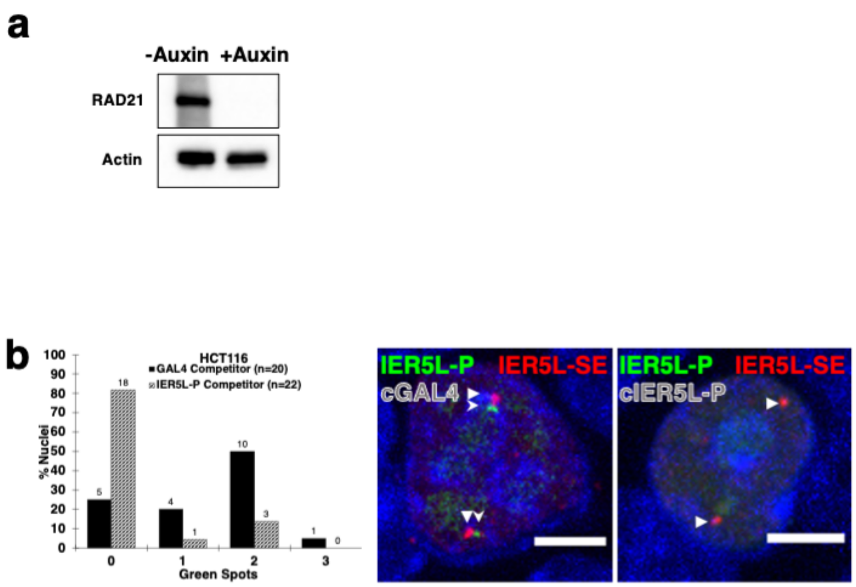


**Supplementary Fig 3. (a)** RAD21 and actin immunoblot of HCT116/RAD21-mAID cell extracts without and with auxin treatment. **(b)** Competitor experiments for validating *IER5L* P-SE labeling. Column plots show percentages of nuclei with the indicated numbers of Clover (green) spots (counts shown on top of columns) in HCT116 cells co-transfected with Clover-PUFc/gIER5L-P-15xPBSc, PUF9R-iRFP670/gIER5L-SE-15xBPS9R and either a non-competing GAL4 gRNA (black columns) or a gRNA competing with the IER5L-P (patterned columns). Representative images in the non-competed or competed samples (Clover, green spots and stealth arrowheads; iRFP670, red spots and triangle arrowheads) are shown to the right. Scale bars, 5 μm.

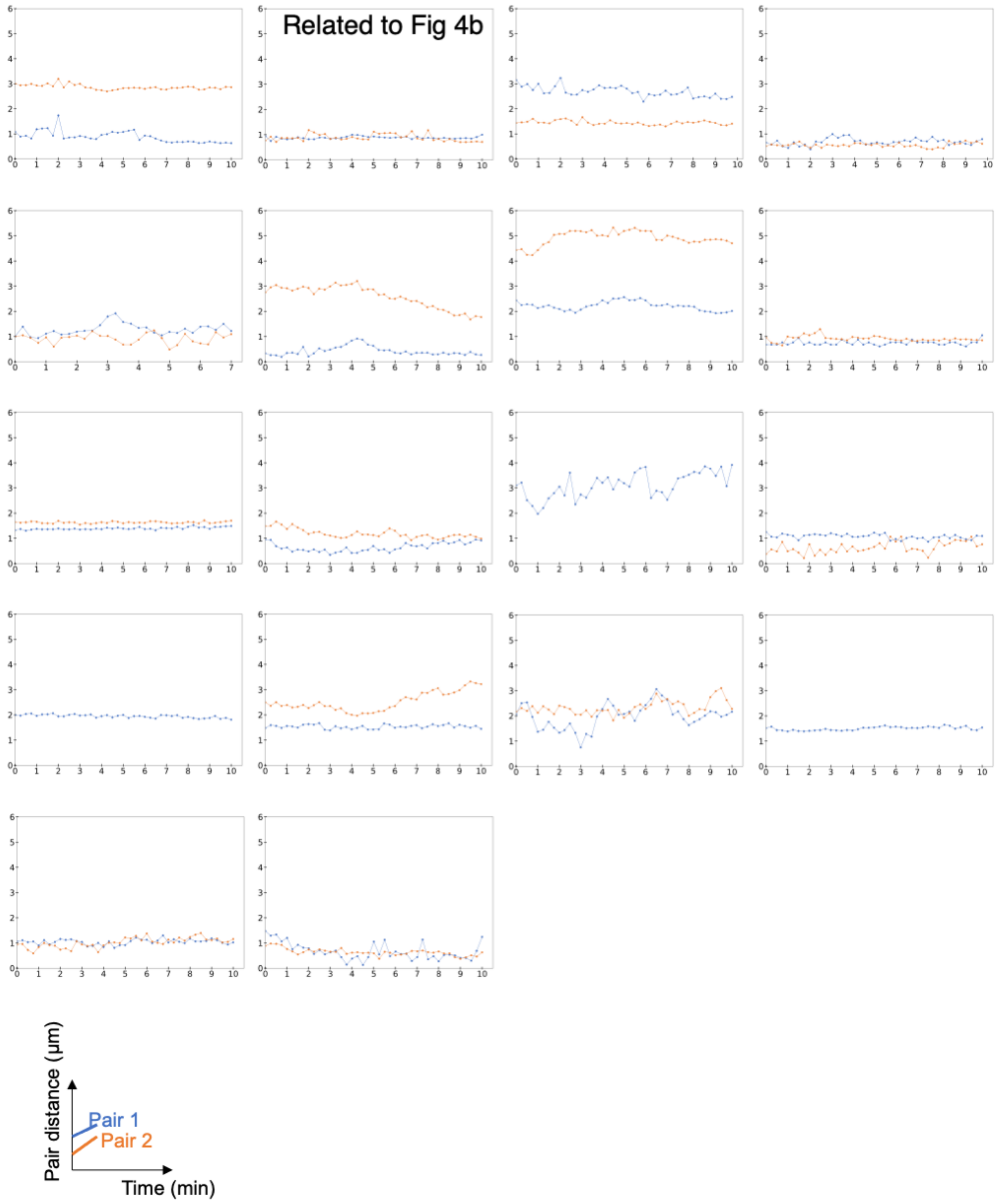


**Supplementary Fig. 4.** Pairwise 3D distances of Clover and iRFP670 foci labeling the *IER5L*-P locus and *IER5L*-SE locus in 18 nuclei (33 pairs) of untreated HCT116/RAD21-mAID cells over time.

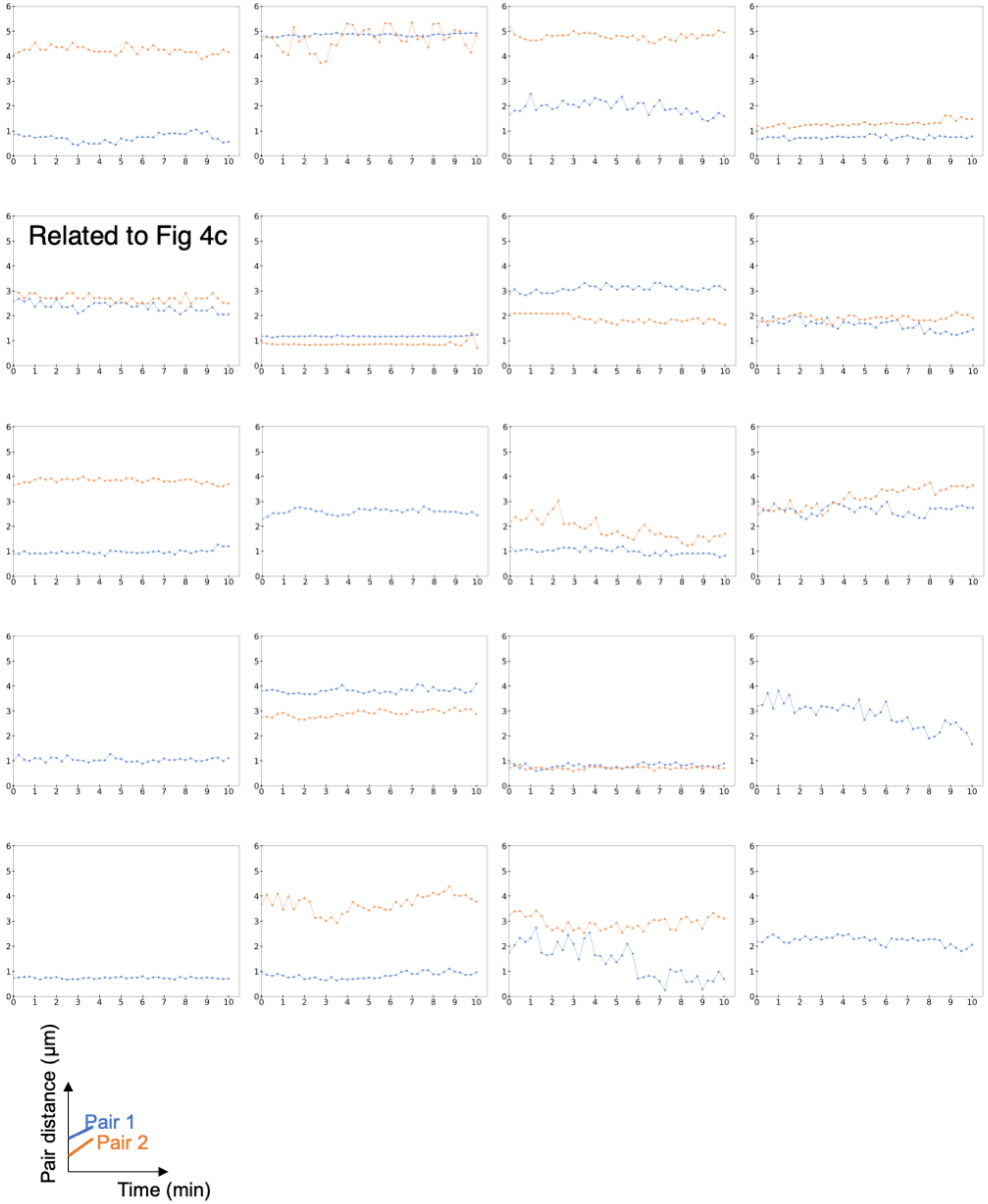


**Supplementary Fig. 5.** Pairwise 3D distances of Clover and iRFP670 foci labeling the *IER5L*-P locus and *IER5L*-SE locus in 20 nuclei (35 pairs) of auxin-treated HCT116/RAD21-mAID cells over time.

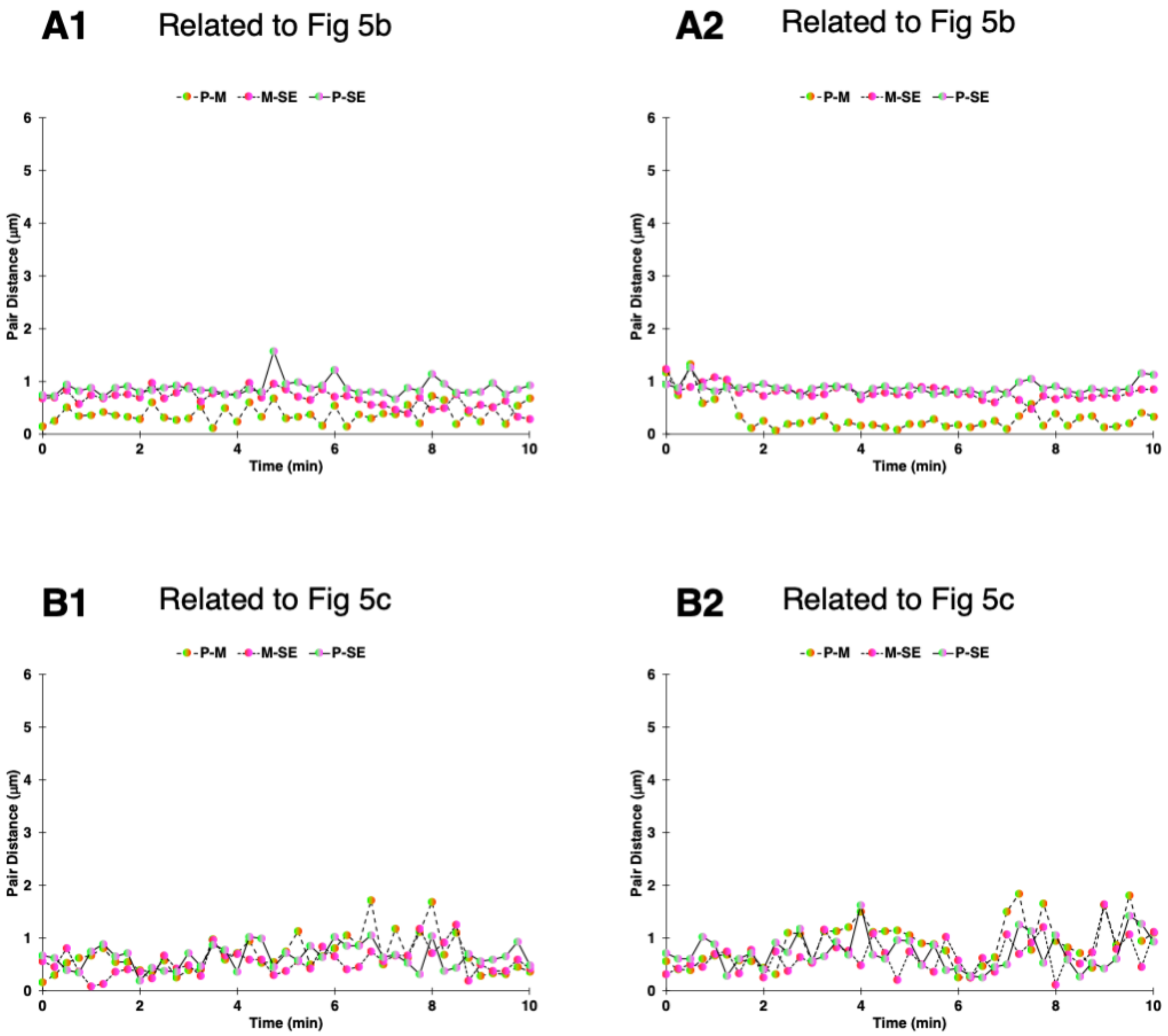


**Supplementary Fig. 6.** All combinations of pairwise 3D distances of Clover (locus A), iRFP670 (locus M) and mRuby2 (locus B) spots on the *IER5L* P-SE loop in 10 HCT116/RAD21-mAID nuclei (15 alleles) over time.

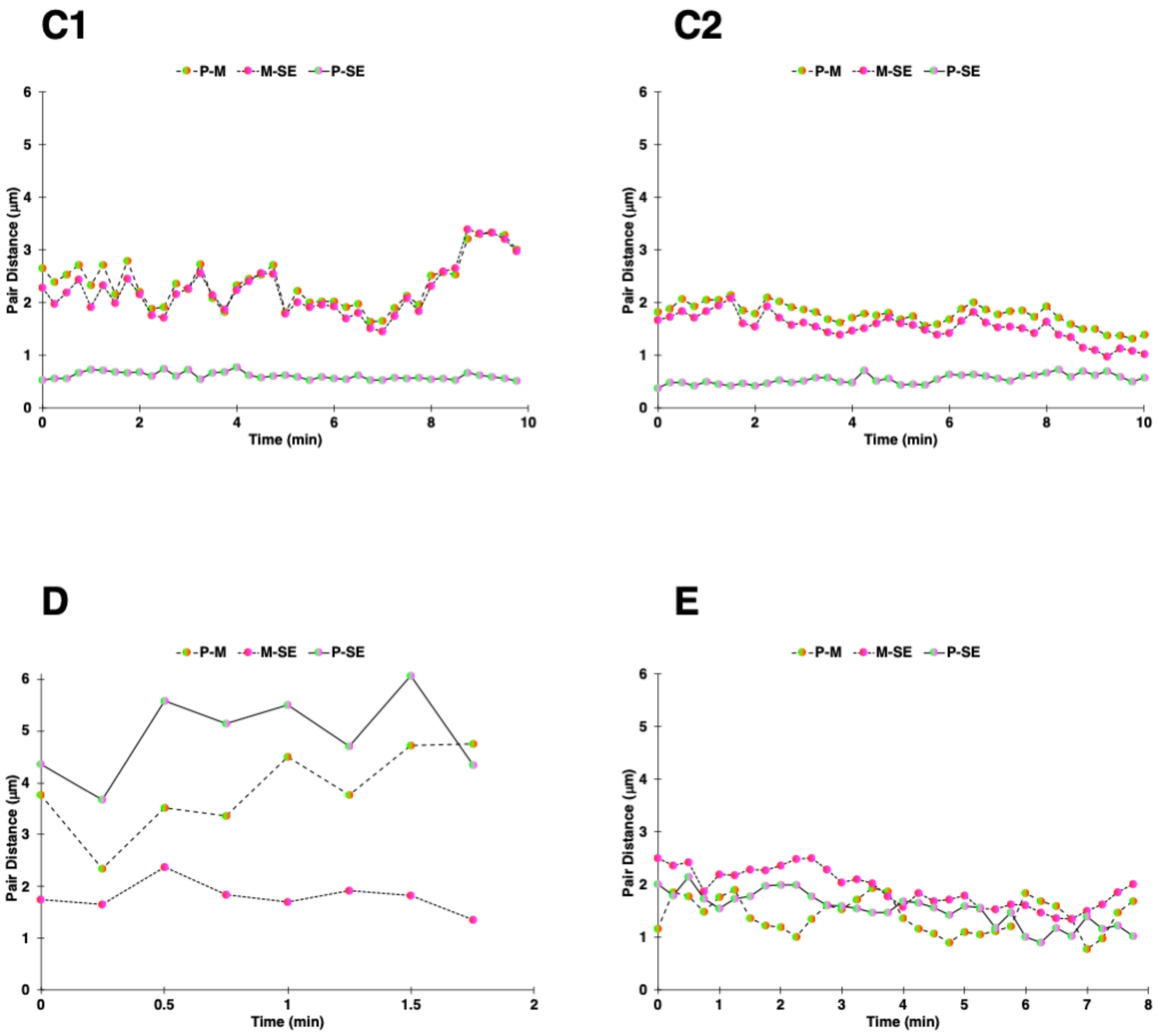


**Supplementary Fig. 6 (continued).** All combinations of pairwise 3D distances of Clover (locus A), iRFP670 (locus M) and mRuby2 (locus B) spots on the *IER5L* P-SE loop in 10 HCT116/RAD21-mAID nuclei (15 alleles) over time.

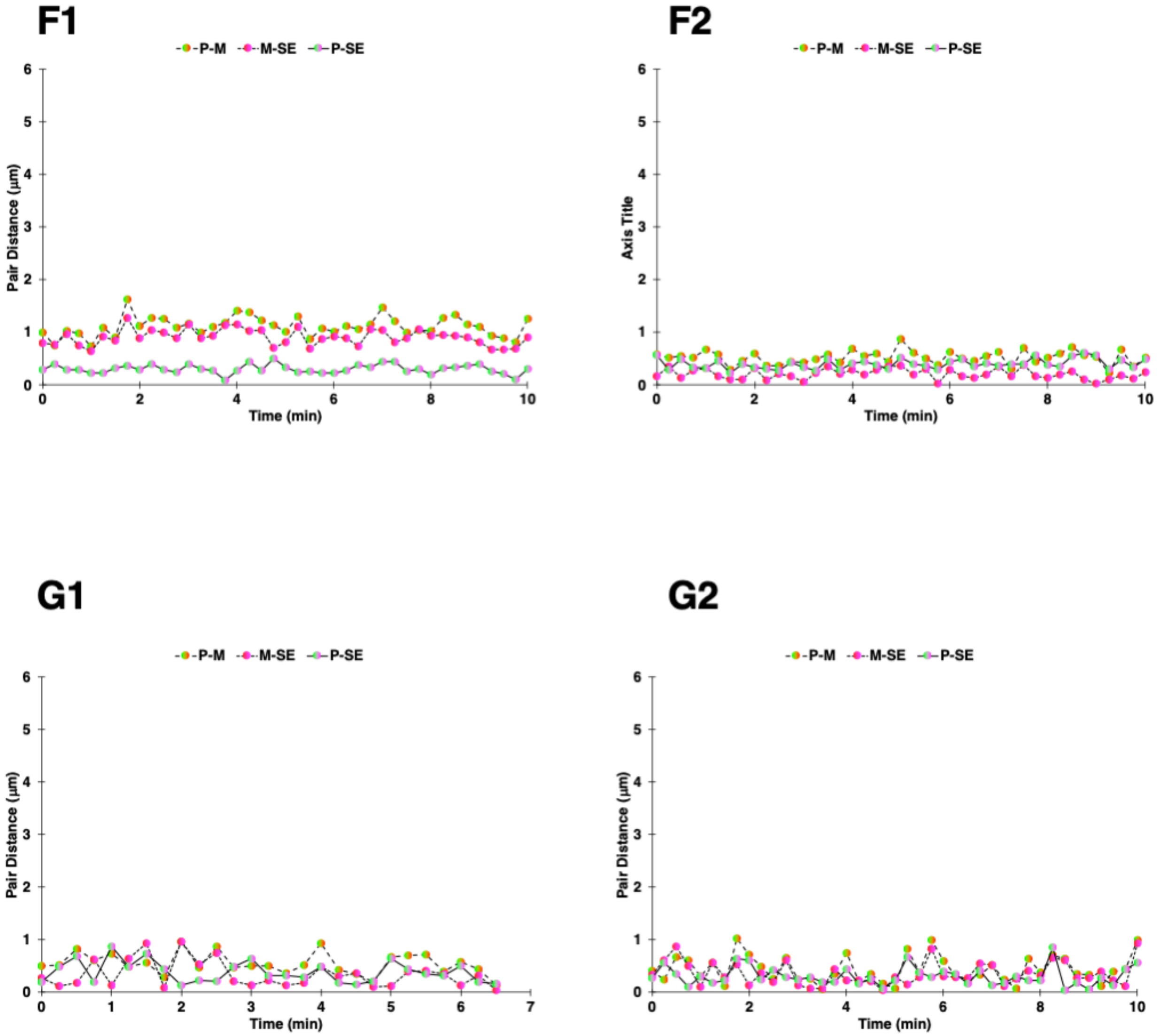


**Supplementary Fig. 6 (continued).** All combinations of pairwise 3D distances of Clover (locus A), iRFP670 (locus M) and mRuby2 (locus B) spots on the *IER5L* P-SE loop in 10 HCT116/RAD21-mAID nuclei (15 alleles) over time.

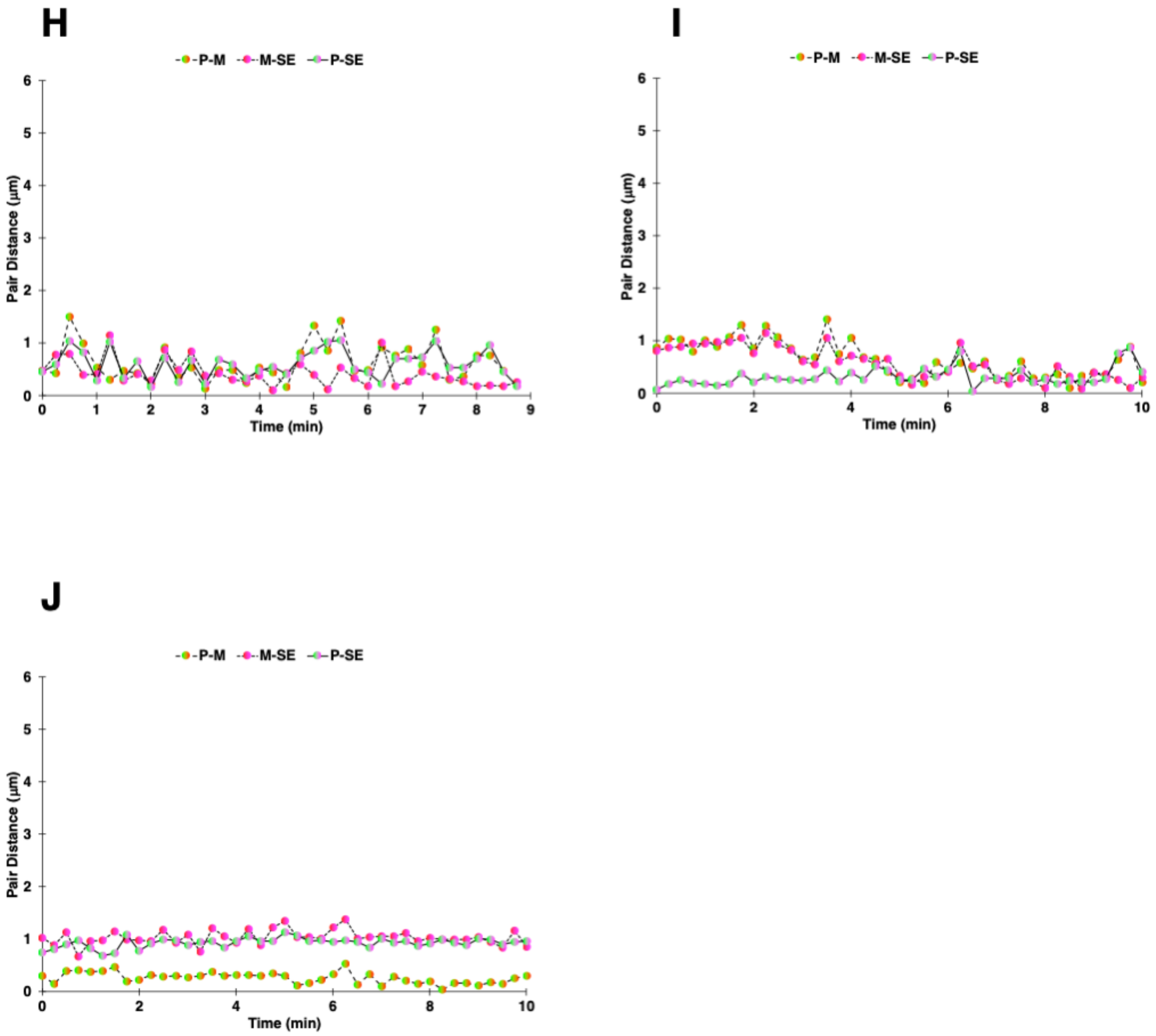


**Supplementary Fig. 6 (continued).** All combinations of pairwise 3D distances of Clover (locus A), iRFP670 (locus M) and mRuby2 (locus B) spots on the *IER5L* P-SE loop in 10 HCT116/RAD21-mAID nuclei (15 alleles) over time.

**Supplementary Video Legends**

**Supplementary Video 1.** Time-lapse video of a representative ARPE-19 nucleus visualized by Casilio labeling the *MASP1-BCL6* loop 5' and 3' anchors with Clover (green) and iRFP670 (red), respectively. Top panels, from left to right: Clover (green), iRFP670 (red), merged (scale bar, 5μm), blue and orange panels correspond to boxed areas of the merged image (scale bars, 1μm). Lower panel shows the pairwise distance (y-axis, μm) with line colors corresponding to the boxed pairs (x-axis, minutes). Related to Fig. 3b.

**Supplementary Video 2.** Time-lapse video of representative untreated HCT116/RAD21-mAID nuclei visualized by Casilio labeling the *IER5L* promoter with Clover (green) and *IER5L* super-enhancer with iRFP670 (red), respectively. Top panels, from left to right: Clover (green), iRFP670 (red), merged (scale bar, 5μm), blue and orange panels correspond to boxed areas of the merged image (scale bars, 1μm). Lower panel shows the pairwise distance (y-axis, μm) with line colors corresponding to the boxed pairs (x-axis, minutes). Related to Fig. 4b.

**Supplementary Video 3.** Time-lapse video of representative auxin-treated HCT116/RAD21-mAID nuclei visualized by Casilio labeling the *IER5L* promoter with Clover (green) and *IER5L* super-enhancer with iRFP670 (red), respectively. Top panels, from left to right: Clover (green), iRFP670 (red), merged (scale bar, 5μm), blue and orange panels correspond to boxed areas of the merged image (scale bars, 1μm). Lower panel shows the pairwise distance (y-axis, μm) with line colors corresponding to the boxed pairs (x-axis, minutes). Related to Fig. 4c.

**Supplementary Video 4.** Time-lapse video of representative untreated HCT116/RAD21-mAID nuclei visualized by Casilio labeling a RAD21-independent loop. Top panels, from left to right: Clover (green), iRFP670 (red), merged (scale bar, 5μm), blue and orange panels correspond to boxed areas of the merged image (scale bars, 1µm). Lower panel shows the pairwise distance (y-axis, µm) with line colors corresponding to the boxed pairs (x-axis, minutes). Related to Fig. 4f.

**Supplementary Video 5.** Time-lapse video of representative auxin-treated HCT116/RAD21-mAID nuclei visualized by Casilio labeling a RAD21-independent loop. Top panels, from left to right: Clover (green), iRFP670 (red), merged (scale bar, 5µm), blue, orange and yellow panels correspond to boxed areas of the merged image (scale bars, 1µm). Lower panel shows the pairwise distance (y-axis, µm) with line colors corresponding to the boxed pairs (x-axis, minutes). Related to Fig. 4g.

**Supplementary Videos 6,7.** PISCES – 3-color 3-point live-cell time-lapse imaging of the *IER5L* promoter-super enhancer loop with promoter, mid-point, super-enhancer labeled by Clover (green, first left panel), iRFP670 (red, second panel), and mRuby2 (magenta, third panel). Right panel shows merged image. Scale bars, 5 µm. Videos 6, 7 related to Fig. 5b, 5c respectively.
